# Variation in genome content and predatory phenotypes between *Bdellovibrio* sp. NC01 isolated from soil and *B. bacteriovorus* type strain HD100

**DOI:** 10.1101/551937

**Authors:** Laura E. Williams, Nicole Cullen, Joseph A. DeGiorgis, Karla J. Martinez, Justina Mellone, Molly Oser, Jing Wang, Ying Zhang

**Author notes:** Corresponding author: Laura E. Williams.

## Abstract

Defining phenotypic and associated genotypic variation among *Bdellovibrio* may further our understanding of how this genus attacks and kills different Gram-negative bacteria. We isolated *Bdellovibrio* sp. NC01 from soil. Analysis of 16S rRNA gene sequences and average amino acid identity showed that NC01 belongs to a different species than the type species *bacteriovorus*. By clustering amino acid sequences from completely sequenced *Bdellovibrio* and comparing the resulting ortholog groups to a previously published analysis, we defined a “core genome” of 778 protein-coding genes and identified four protein-coding genes that appeared to be missing only in NC01. To determine how horizontal gene transfer (HGT) may have impacted NC01 genome evolution, we performed genome-wide comparisons of *Bdellovibrio* nucleotide sequences, which indicated that eight NC01 genomic regions were likely acquired by HGT. To investigate how genome variation may impact predation, we compared protein-coding gene content between NC01 and the *B. bacteriovorus* type strain HD100, focusing on genes implicated as important in successful killing of prey. Of these, NC01 is missing ten genes that may play roles in lytic activity during predation. Compared to HD100, NC01 kills fewer tested prey strains and kills *E. coli* ML35 less efficiently. NC01 causes a smaller log reduction in ML35, after which the prey population recovers and the NC01 population decreases. In addition, NC01 forms turbid plaques on lawns of *E. coli* ML35, in contrast to clear plaques formed by HD100. Linking phenotypic variation in interactions between *Bdellovibrio* and Gram-negative bacteria with underlying *Bdellovibrio* genome variation is valuable for understanding the ecological significance of predatory bacteria and evaluating their effectiveness in clinical applications.

## Introduction

Since their discovery in the early 1960s, predatory bacteria have been isolated from a wide range of terrestrial and aquatic environments, including some with extreme conditions such as high temperature or salinity (1). The impact of predatory bacteria in these environments is not fully known, but recent studies indicate they may play an important role in shaping microbial communities (2–4). In addition, their ability to attack and kill Gram-negative bacterial pathogens has generated interest in developing predatory bacteria as biocontrol agents to combat drug-resistant infections (5–7). Given their potential ecological significance and possible clinical applications, it is important to define phenotypic and genotypic variation among predatory bacteria. In particular, variation in predatory phenotypes such as prey range and predation efficiency will affect how different strains of predatory bacteria impact microbial communities, and this phenotypic variation may be determined by variation in genome content. To further our understanding of phenotypic and genome variation in predatory bacteria, we isolated, assayed and sequenced a *Bdellovibrio* strain from soil and compared its predatory phenotypes and genome content to other sequenced *Bdellovibrio*.

Among the different genera of predatory bacteria, *Bdellovibrio* is the most well-studied. Currently, this genus includes two species that differ in their predation strategy. *B. bacteriovorus* invades the periplasm of Gram-negative prey cells (8), whereas *B. exovorus* attaches to the outside of prey cells but does not enter the cell itself (9). Here, we focus on intraperiplasmic *Bdellovibrio*. This lifecycle includes two main phases: the attack phase and the growth phase (10). During the attack phase, small, highly motile *Bdellovibrio* cells seek out and attach to Gram-negative prey bacteria. *Bdellovibrio* then invades the prey periplasm by using lytic enzymes to open a small hole in the prey outer membrane and cell wall (11). Both attachment and invasion may be mediated by Type IV pili expressed by *Bdellovibrio* (12–14). After *Bdellovibrio* moves into the prey periplasm, it uses peptidoglycan-modifying enzymes to reseal and reshape the prey cell wall, resulting in formation of a rounded structure called a bdelloplast (15). Within the bdelloplast, *Bdellovibrio* initiates the growth phase, during which it secretes lytic enzymes into the prey cytoplasm. Using digested prey cell contents as nutrients, *Bdellovibrio* grows into a single long filament within the prey periplasm, eventually dividing into multiple progeny, which lyse the bdelloplast and emerge as new attack phase cells.

By invading and lysing Gram-negative bacteria, some strains of *Bdellovibrio* can reduce prey populations by up to 8 log units over 48 hours *in vitro* (6, 16). However, not all *Bdellovibrio* strains are equally effective at killing the same Gram-negative species. Researchers have observed variation among *Bdellovibrio* strains in which Gram-negative bacteria are susceptible to predation (17, 18) and in how quickly and to what extent a population of Gram-negative prey is reduced (16, 19). These predatory phenotypes, referred to as prey range and predation efficiency, are clearly important in determining the outcomes of interactions between *Bdellovibrio* and Gram-negative bacteria; however, the extent of naturally occurring variation in these phenotypes and the underlying mechanisms governing them are not well understood. Defining how predatory phenotypes differ among *Bdellovibrio* will contribute to our understanding of the impact of predatory bacteria on microbial communities in different environments and inform their application in biocontrol of bacterial pathogens.

Here, we describe the isolation of *Bdellovibrio* sp. NC01 from soil and compare its genome and predatory phenotypes to other *Bdellovibrio*, focusing primarily on the *B. bacteriovorus* type strain HD100. The intraperiplasmic predator HD100 was the first *Bdellovibrio* strain sequenced (20), and multiple studies have dissected some of the molecular mechanisms governing predation in this strain (21). To investigate variation in genome content, we compared the genome of NC01 to that of HD100 and other sequenced *Bdellovibrio*. Currently, only a few completely sequenced predatory bacteria genomes are available in public databases. In addition to HD100, there are only four other completely sequenced intraperiplasmic *Bdellovibrio* genomes, including *B. bacteriovorus* W (22), which is the most phylogenetically divergent from HD100 (23). The limited genome dataset highlights the importance of sequencing additional *Bdellovibrio*, especially strains that are divergent from HD100. To define variation in predatory phenotypes between NC01 and HD100, we determined prey range differences by testing each strain against a panel of Gram-negative bacteria, and we measured predation efficiency by quantifying viable *E. coli* ML35 during co-culture with each strain.

## Methods

### Isolation and identification of environmental bacteria for use as prey

We isolated bacteria from an urban freshwater stream in Providence, RI (lat 41.835, lon - 71.44299) and from soil in a residential area of Massachusetts (lat 42.0877, lon −71.2309) for use as prey. We also obtained *E. coli* ML35 from Mark Martin (University of Puget Sound, Puget Sound, WA) and *Escherichia* 0057 from Brett Pellock (Providence College, Providence, RI).

To identify these isolates, we sequenced the 16S rRNA gene following a previously described procedure (24). Briefly, we used primers 63F (25) and 1378R (26), which amplify the 16S rRNA gene from many members of Domain Bacteria. Sanger sequencing of purified PCR products was performed by GeneWiz (South Plainfield, NJ) using the same primers. We trimmed and assembled reads using Phred/Phrap/Consed (27–29), then classified the assembled 16S rRNA gene sequences using BLAST (30), the SILVA Incremental Aligner (31) and the Ribosomal Database Project classifier (32). Supplementary Table 1 shows the full classification results. The RDP classifier and the SILVA Incremental Aligner classify sequences to the genus level, but not the species level; therefore, species names are not provided for these isolates.

### Isolation and identification of *Bdellovibrio* sp. NC01 from bioswale soil

In September 2015, we collected 50 g of soil from a bioswale on the Providence College campus (lat 41.84277, lon −71.43944). To isolate predatory bacteria from the soil sample, we adapted standard methods (33, 34). We combined the soil sample with 500 ml of sterile HM buffer (25 mM HEPES adjusted to pH 7.4 and supplemented with 3 mM calcium chloride dihydrate and 2 mM magnesium chloride hexahydrate). We then filtered this mixture through sterile cheesecloth into a sterile container to remove large particulate matter. We vortexed the filtrate to release bacterial cells from the small particles of soil that passed through the cheesecloth. To pellet the soil particles, we centrifuged the filtrate at 500 × g (1753 rpm in a Sorvall GSA rotor) for 10 minutes, after which we retained the supernatant containing suspended bacterial cells.

To enrich for predatory bacteria in the soil sample supernatant, we used *Pseudomonas* sp. NC02 as prey, because it was also isolated from a soil environment, and its genome has been sequenced (35). We cultured *Pseudomonas* sp. NC02 overnight in 120 ml tryptic soy broth (TSB) at 28°C and 200 rpm, then centrifuged the culture at 5854 × g (6000 rpm in a Sorvall GSA rotor) for 10 minutes, washed the resulting pellet with 100 ml sterile HM buffer, centrifuged again and resuspended the resulting pellet in 2.5 ml HM buffer. We combined 1 ml of *Pseudomonas* sp. NC02 resuspension with 20 ml of soil sample supernatant and incubated this enrichment at 28°C and 200 rpm.

Once we observed small, fast-moving cells under 1000x phase-contrast microscopy, we filtered the enrichment using a 0.45 μm filter and performed 10-fold serial dilutions of the filtrate in sterile HM buffer. Dilutions were then plated using a double agar overlay method. For each dilution of the filtrate to be plated, we combined 100 μl of the dilution, 250 μl of *Pseudomonas* sp. NC02 at 10^8^ cfu/ml and 4.5 ml of molten HM top agar (0.6% agar). After vortexing, we poured this mixture onto HM agar plates (1.5% agar). We allowed the top agar to solidify at room temperature, then incubated plates at 28°C until we observed plaques in the lawn of *Pseudomonas* sp. NC02. To begin the process of obtaining a pure isolate, we used a sterile Pasteur pipette to extract single, well-defined plaques from the top agar. We combined each plaque with 500 μl of sterile HM buffer, incubated the plaque suspension at room temperature for at least 5 minutes, then vortexed it vigorously.

To culture predatory bacteria from the plaque suspensions, we made lysates combining 300 μl of each plaque suspension, 1.5 ml of an overnight culture of *Serratia* 0043 and 20 ml sterile HM buffer. *Serratia* 0043 was also isolated from a soil environment, and switching to this prey species during the isolation procedure selected for predatory isolates that were capable of attacking both *Pseudomonas* and *Serratia*, which belong to different orders. We incubated lysates at 28°C and 200 rpm. After 48 h, one lysate showed small, fast-moving cells, and we used this lysate to begin two additional rounds of serial dilution, plating and plaque picking to ensure a pure isolate. We used *Serratia* 0043 as prey during both of these rounds. After the final round, we filtered the plaque suspension using a 0.45 μm filter, then combined 500 μl of filtrate with 500 μl of sterile 50% glycerol and stored the stock at −80°C.

To identify NC01, we extracted the 16S rRNA gene sequence from the NC01 genome based on the annotation generated by PGAP (see below for genome sequencing and annotation methods). Using SINA 1.2.11 (31), we aligned this sequence against 16S rRNA gene sequences extracted from nine genomes of obligate predatory bacteria belonging to *Bacteriovorax*, *Bdellovibrio* and *Halobacteriovorax* (*Bacteriovorax stolpii* DSM12778 CP025704, *Bdellovibrio bacteriovorus* 109J CP007656, *Bdellovibrio bacteriovorus* HD100 NC_005363, *Bdellovibrio bacteriovorus* SSB218315 CP020946, *Bdellovibrio bacteriovorus* Tiberius NC_019567, *Bdellovibrio bacteriovorus* W NZ_CP002190, *Bdellovibrio exovorus* JSS NC_020813, *Halobacteriovorax marinus* BE01 CP017414, *Halobacteriovorax marinus* SJ NC_016620). We then inferred a phylogenetic tree from the alignment with the online server RAxML Blackbox (36), using 100 bootstrap replicates to estimate the confidence of the topology.

To further investigate the relationships among NC01 and other *Bdellovibrio*, we used an average amino acid identity (AAI) calculator (http://enve-omics.ce.gatech.edu/) (37) to analyze the similarity among protein-coding sequences from NC01, 109J, HD100 and W, as annotated by RAST (see below for genome sequencing and annotation methods).

### Negative stain electron microscopy of *Bdellovibrio* sp. NC01

To obtain samples of *Bdellovibrio* sp. NC01 for negative stain electron microscopy, we added a small amount of the −80°C freezer stock to 20 ml of HM buffer mixed with 1.5 ml of an overnight culture of *E. coli* ML35 grown in TSB. After 48 h of incubation at 28°C and 200 rpm, we stained samples with 1% uranyl acetate (pH 4.2) following a previously described procedure (24) and imaged the resulting specimens using a JEOL CX 2000 transmission electron microscope.

### Genome sequencing and annotation of *Bdellovibrio* sp. NC01

To obtain genomic DNA, we combined a small amount of the −80°C stock, 1.5 ml of an overnight culture of *E. coli* ML35 and 25 ml of sterile HM buffer. We chose ML35 as prey because it is a standard prey strain used in cultivation of *Bdellovibrio*, and the genome sequence is available for screening reads if needed (38). After 72 hours of incubation at 28°C and 200 rpm, we centrifuged the lysate at 10000 × g (9100 rpm in a Thermo Scientific A27 rotor) for 10 minutes to pellet bacterial cells. Because PacBio sequencing technology requires at least 10 μg of genomic DNA, we did not filter the lysate to prevent loss of predatory bacteria cells. We then used the Wizard Genomic DNA purification kit to extract genomic DNA, following the manufacturer’s instructions beginning with resuspension of the pellet in 600 μl of the kit’s nuclei lysis solution. For the final rehydration step, we rehydrated the DNA pellet overnight at 4°C. Quantification by Qubit estimated a final concentration of 340 ng/μl genomic DNA.

For long-read sequencing, library construction and sequencing were performed at the Institute for Genome Sciences at the University of Maryland Baltimore on a Pacific Biosciences RSII instrument using P6-C4 chemistry. Quality control analysis of the reads showed that <10% of reads aligned by BLAST to *E. coli* or *Shigella*, indicating that representation of the prey genome in the read dataset is low. To analyze and assemble the reads, we launched an instance of SMRT Portal 2.3.0 using Amazon EC2. A single SMRT cell yielded 79,088 postfilter polymerase reads (N_50_ 17,028) and 94,031 subreads (N_50_ 13,572). Within SMRT Portal, we assembled the reads with HGAP3 (39), using 4.5 Mbp for the genome size. This generated 186 contigs. The largest contig was 4,001,137 bp and aligned to *Bdellovibrio* in the database using BLASTN. The other 185 contigs were each <48,000 bp in length, and all of them aligned to *E. coli* by BLASTN, which is expected due to the inclusion of prey genomic DNA in the sequencing library.

To close the genome, we used BLASTN to identify overlap in the ends of the contig and trimmed the sequence to generate a closed contig of 3,975,771 bp. We also adjusted the start site so that the first nucleotide of the closed genome sequence corresponds to the first nucleotide in the start codon of *dnaA*. To polish this sequence, we used 250 bp paired-end Illumina MiSeq data generated at the Rhode Island Genomics and Sequencing Center. We filtered and trimmed the raw read dataset to retain only those read pairs for which the quality score of every base was ≥13 and the read length was ≥65 nt. This yielded 1,256,135 read pairs. Using bwa-mem (40), samtools (41) and Pilon (42), we aligned these reads to the closed genome sequence and corrected 1 SNP, 84 small insertions totaling 85 bases and 4 small single-base deletions. We manually examined changes flagged by Pilon using the alignment viewer Tablet (43), generating a final, polished *Bdellovibrio* sp. NC01 genome sequence of 3,975,853 bp. This sequence has been deposited in GenBank as CP030034.

In addition to annotation with NCBI’s Prokaryotic Genome Annotation Pipeline (PGAP) version 4.5, we also annotated the sequence using ClassicRAST (44, 45), Infernal (46), and tRNAScan-SE (47, 48).

### Comparative genomics of *Bdellovibrio* sp. NC01 and other sequenced *Bdellovibrio*

We used a custom software pipeline (49) to cluster amino acid sequences from *Bdellovibrio* sp. NC01, all five completely sequenced *B. bacteriovorus* genomes, and *B. exovorus* JSS into ortholog groups. NCBI accession numbers for these genomes are listed above. Ortholog groups were identified among all genomes based on the identification of bi-directional best hits in the protein-coding genes using BLASTP for the pairwise sequence comparisons. The pipeline requires an alignment of two amino acid sequences to have ≥30% identity, ≥70% coverage, and an e-value of <0.001. To define the “core genome” of the seven *Bdellovibrio* analyzed here, we identified the set of ortholog groups that each included at least one protein-coding gene from each genome.

We compared the core genome defined by our analysis with the set of 795 conserved *Bdellovibrio* genes identified in (23). To do this, we extracted NCBI protein accession numbers associated with *B. exovorus* JSS from Additional File 4a, because JSS was the only genome for which all 795 genes were listed. We then used R and bash scripts to determine whether these accession numbers appeared in the ortholog groups identified as the core genome in our analysis.

To define the set of HD100 protein-coding genes implicated in predation by previous studies, we extracted the HD100 locus tags listed in Supporting Information Figure S1 of (50) and all HD100 locus tags with a value <0.5 in the column labelled Mini Tn-seq ECPL avg *W* in Data Set S4 of (51). For each of these HD100 locus tags, we used R and bash scripts to determine the corresponding GenBank protein ID (as listed in the NC_005363 annotation on GenBank) and identify the ortholog group in our analysis that included this protein ID. We then determined whether the ortholog group also included a NC01 protein-coding gene. If not, we considered the gene missing in NC01.

For genome-wide pairwise comparisons of nucleotide sequences, we used the BLAST Ring Image Generator (BRIG) (52).

### Phenotype assays to test prey range differences and predation efficiency

We obtained *Bdellovibrio bacteriovorus* HD100 from Mark Martin (University of Puget Sound, Puget Sound, WA) for predatory phenotype comparisons. To conduct phenotype assays, we adapted standard methods (33, 34, 53). To culture *Bdellovibrio* NC01 and HD100, we added a small amount of −80°C stocks to 20 ml HM buffer combined with 1.5 ml of an overnight culture of *E. coli* ML35. After 72 hours of incubation at 28°C and 200 rpm, we filtered the lysate using a 0.45 μm filter to separate predatory cells from prey cells and cell debris. For each phenotype assay described below, we followed this protocol to obtain attack phase *Bdellovibrio* cells.

To test prey range differences and plaque appearance, we plated 10-fold serial dilutions of filtered lysate using a double agar overlay method as described above for the isolation procedure. Each tested prey strain was grown in 120 ml TSB overnight at 28°C and 200 rpm, with the exception of *Escherichia* strains, which were grown at 37°C. After centrifugation and washing, we resuspended prey cell pellets in 5 ml of sterile HM buffer. Based on cfu/ml counts, this yielded concentrations of at least 10^8^ cfu/ml, which ensured a dense prey lawn for visualizing plaques. For some prey strains, we added 500 μl of prey resuspension to top agar to improve our ability to detect and observe plaques in the prey lawn. We incubated plates at 28°C for 8 days before scoring plaque formation and performed three replicates for each prey strain. In observations of plaque appearance, we photographed plates at multiple time points over the course of 8 days.

To test predation efficiency, we adapted a protocol described in (19). We centrifuged filtered lysate at 8635 × g (8500 rpm in a Thermo Scientific A27 rotor) for 10 minutes. The resulting pellet of predatory bacteria was resuspended in sterile HM buffer to yield ~10^8^ cells/ml based on direct cell counts using a Petroff-Hausser counting chamber and 1000x phase-contrast microscopy. To initiate the predation efficiency assay, we combined 500 μl of the *Bdellovibrio* cell resuspension with 12.5 ml of an overnight culture of *E. coli* ML35 diluted to OD_600_ 0.15 (range: 0.135 – 0.165) in sterile HM buffer. Dilution of the overnight culture yielded ~10^8^ *E. coli* cells/ml based on cfu/ml counts. As a control, we used 500 μl of sterile HM buffer in place of *Bdellovibrio* cell resuspension. Using cfu/ml counts, we quantified viable *E. coli* ML35 at the beginning of the assay (time zero) and then after 6, 24, 48 and 72 hours of incubation at 28°C and 200 rpm. Based on the initial concentrations of *Bdellovibrio* and *E. <u>coli</u>* as estimated by direct cell counts and cfu/ml counts, the predator:prey ratio at the beginning of the assay was 1:5 for NC01 and 1:4 for HD100.

To test survival of *Bdellovibrio* in the absence of prey, we used the same protocol as the predation efficiency assay, but we combined 500 μl of *Bdellovibrio* cell resuspension with 12.5 ml of sterile HM buffer instead of a dilution of prey overnight culture. After 0, 24, 48 and 72 hours of incubation at 28°C and 200 rpm, we quantified viable *Bdellovibrio* by plating serial dilutions of flask contents using the double agar overlay method described above for testing prey range differences. After incubating these plates for 48 hours at 28°C, we counted plaques formed on the *E. coli* ML35 prey lawn and then obtained a final plaque count after another 24 hours of incubation.

Because we used both direct cell counts and plaque counts to quantify *Bdellovibrio*, we validated the methods by comparing initial *Bdellovibrio* concentrations for the survival assay without prey as estimated by direct cell counts using the Petroff-Hausser counting chamber and plaque counts using serial dilution and double agar overlay. If all cells counted in the Petroff-Hausser counting chamber are viable, these estimates should be the same. For both NC01 (n=4) and HD100 (n=3), the two quantification methods showed good agreement (Supplementary Figure 1). The average initial concentration estimated by direct cell counts was 1.05 × 10^7^ cells/ml for NC01 and 2.3 × 10^7^ cells/ml for HD100, whereas the average initial concentration estimated by plaque counts (reported as plaque-forming units (pfu)/ml) at time zero was 1.17 × 10^7^ pfu/ml for NC01 and 2.71 × 10^7^ pfu/ml for HD100. These data show that direct cell counts and plaque counts provide comparable results for quantification of *Bdellovibrio*.

## Results

### Bioswale soil harbors predatory bacteria belonging to *Bdellovibrio*

Using Gram-negative soil bacteria isolated from a residential area as prey, we performed an enrichment followed by three rounds of plaque picking to obtain a pure isolate of predatory bacteria from a sample of soil collected at a bioswale on the Providence College campus. The bioswale is a manmade landscape feature designed to collect and filter stormwater runoff from nearby buildings. We designated the bioswale soil isolate as NC01. Using 1000x phase-contrast microscopy of lysates combining NC01 and Gram-negative prey, we observed small, fast-moving attack phase NC01 cells that attached to prey cells (Figure 1a). We also observed rounded structures characteristic of bdelloplasts (Figure 1b), suggesting that NC01 uses a cell invasion strategy similar to *Bdellovibrio bacteriovorus*. Negative stain electron microscopy showed that NC01 attack phase cells have a comma-shaped morphology with a single polar flagellum (Figure 1c).

**Figure 1.**
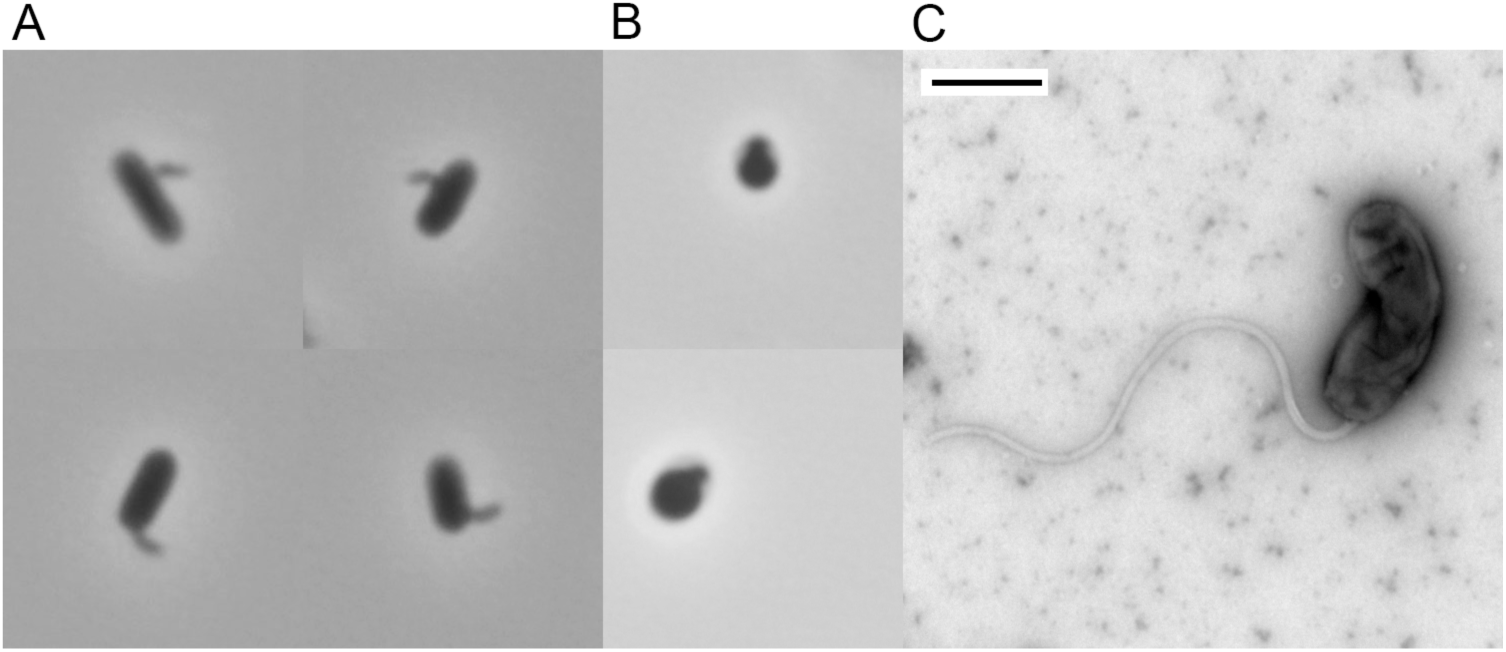
Microscopy of *Bdellovibrio* sp. NC01 isolated from bioswale soil. (A) Compilation of four images taken using 1000x phase-contrast microscopy to show small, comma-shaped NC01 attack phase cells attached to larger *E. coli* ML35 cells after 48 hours of co-culture. (B) Compilation of two images taken using 1000x phase-contrast microscopy to show rounded structures characteristic of bdelloplasts after 48 hours of co-culture. (C) Negative stain electron microscopy of a NC01 attack phase cell. The scale bar is 500 nm.

To identify NC01, we constructed a phylogenetic tree of 16S rRNA gene sequences from this isolate, all five *Bdellovibrio bacteriovorus* with complete genomes available in GenBank, and representatives of *Bacteriovorax*, *Bdellovibrio exovorus* and *Halobacteriovorax* with complete genomes available in GenBank (Figure 2). In this phylogenetic tree, NC01 clearly clusters with other *Bdellovibrio*. Pairwise alignment of 16S rRNA gene sequences from NC01 and *Bdellovibrio bacteriovorus* HD100, which is the type strain of the species, showed 97% similarity across 1,516 nucleotides. This value is above the 95% similarity cutoff typically used to define bacterial genera (54), which supports identifying NC01 as *Bdellovibrio*. However, it is below the 98.7% similarity cutoff typically used to define species (55), which suggests that NC01 belongs to a different species than *bacteriovorus*.

**Figure 2.**
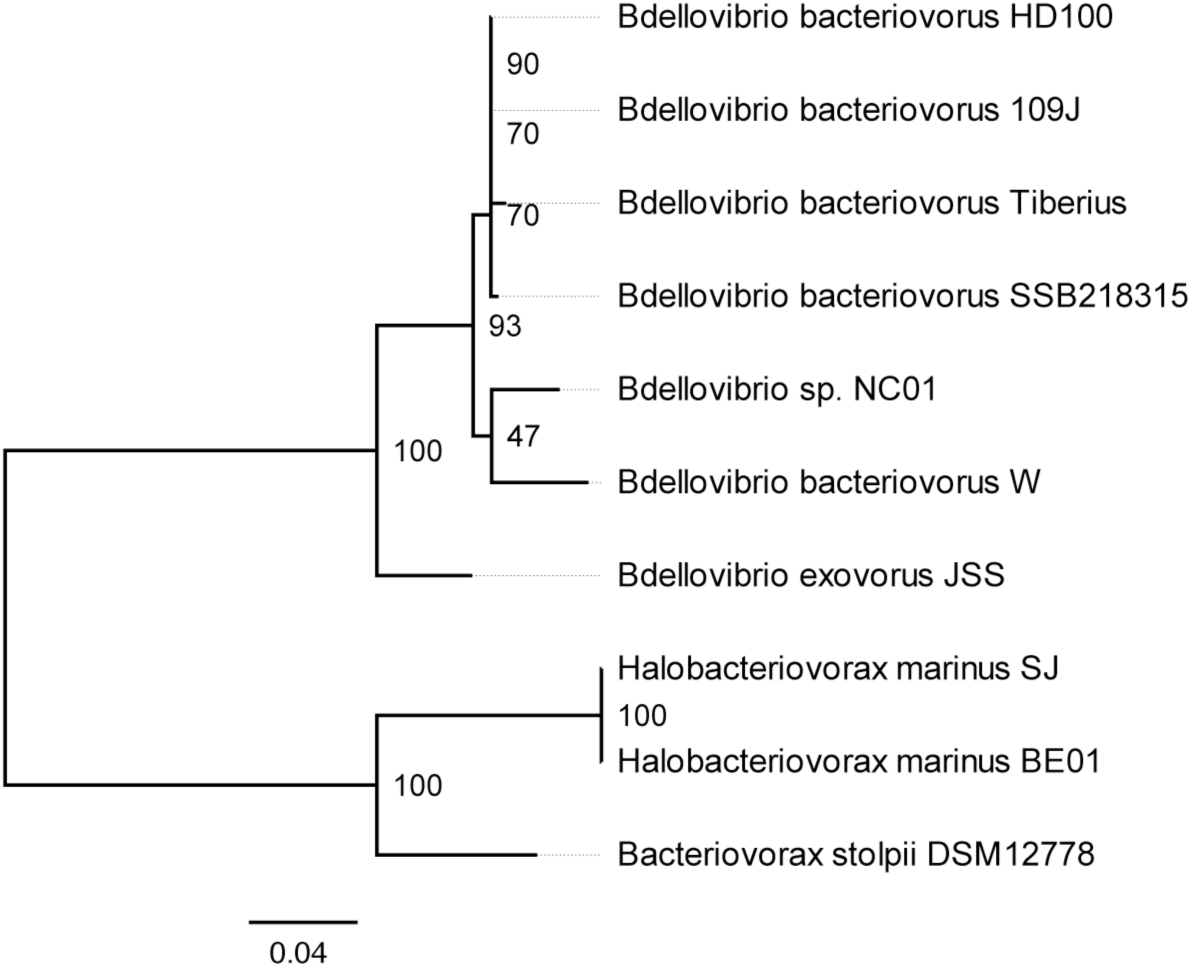
16S rRNA gene phylogenetic tree. We aligned 16S rRNA gene sequences using SINA 1.2.11, then constructed a phylogenetic tree using RAxML BlackBox with 100 bootstrap replicates. The phylogenetic tree is rooted on the branch joining *Bdellovibrio* with *Bacteriovorax*/*Halobacteriovorax*, and bootstrap support for each node is shown.

To explore this further, we compared the average amino acid identity (AAI) of *Bdellovibrio* strains NC01, W, HD100 and 109J (Table 1). HD100 and 109J, which are both identified as *B. bacteriovorus*, cluster in a well-supported clade in the 16S rRNA phylogenetic tree, and their amino acid sequences are highly similar (average AAI 99.2%). However, NC01 amino acid sequences are <70% similar on average to HD100 and 109J amino acid sequences, supporting the conclusion that NC01 belongs to a different species than these two strains. NC01 amino acid sequences are even less similar on average to those of strain W, suggesting that NC01 does not belong to the same species as this strain. Based on these analyses, we chose not to designate a species for NC01, and we refer to this isolate as *Bdellovibrio* sp. NC01.

**Table 1.** Average amino acid identity among representative *Bdellovibrio* genomes

|  | NC01 | W | HD100 | 109J |
| --- | --- | --- | --- | --- |
| NC01 |  | 64.4 | 68.8 | 68.8 |
| W | 64.4 |  | 66.4 | 66.5 |
| HD100 | 68.8 | 66.4 |  | 99.2 |
| 109J | 68.8 | 66.5 | 99.2 |  |

### Analysis of ortholog groups identifies protein-coding genes that are missing in *Bdellovibrio* sp. NC01 but conserved in other sequenced *Bdellovibrio*

The genome of *Bdellovibrio* sp. NC01 is larger than that of both HD100 and W (Table 2), with a lower GC content than that of HD100, but slightly higher than that of W. We compared the NC01 genome to all five publicly available complete *Bdellovibrio bacteriovorus* genomes and the *Bdellovibrio exovorus* JSS genome using a software pipeline that clusters protein-coding genes into ortholog groups based on amino acid sequence similarity (49). Because the RAST and PGAP annotation tools generate slightly different annotations of the same genome sequence, we performed separate analyses of ortholog groups using either RAST or PGAP annotations of the seven genomes in our dataset. To determine the “core genome” in each analysis, we identified ortholog groups that included at least one protein-coding gene from each genome. The core genome based on RAST annotations included 1,364 ortholog groups, whereas the core genome based on PGAP annotations included 1,394 ortholog groups.

**Table 2.** Genome comparison of *Bdellovibrio* sp. NC01, *Bdellovibrio bacteriovorus* HD100 and *Bdellovibrio bacteriovorus* W

|  | *Bdellovibrio*<br>sp. NC01 |  | *Bdellovibrio*<br>*bacteriovorus* HD100 |  | *Bdellovibrio*<br>*bacteriovorus* W |  |
| --- | --- | --- | --- | --- | --- | --- |
|  | PGAP | RAST and<br>RFAM/<br>tRNAScan | PGAP | RAST and<br>RFAM/<br>tRNAScan | PGAP | RAST and<br>RFAM/<br>tRNAScan |
| Genome size (bp) | 3,975,853 |  | 3,782,950 |  | 3,006,992 |  |
| GC content (%) | 44.5 |  | 50.6 |  | 43.3 |  |
| Genes | 3,840 | 3,828 | 3,597 | 3,651 | 2,834 | 2,901 |
| Protein-coding | 3,773 | 3,776 | 3,548 | 3,597 | 2,760 | 2,849 |
| Hypothetical<br>proteins | 1,712 | 1,431 | 1,245 | 1,309 | 962 | 971 |
| tRNA | 34 | 34 | 36 | 36 | 34 | 34 |
| rRNA | 6 | 6 | 6 | 6 | 6 | 6 |
| Other RNA | 4 | 12 | 4 | 12 | 4 | 12 |

Recently, Oyedara and colleagues conducted comparative genomics of the same six publicly available complete *Bdellovibrio* genomes and the genome of *Bdellovibrio* sp. SKB1291214, which is not complete (23). The core genome shared by these seven *Bdellovibrio* strains included 795 protein-coding genes. We compared these to the PGAP core genome described above and identified 778 protein-coding genes that were conserved in all eight *Bdellovibrio* genomes. Of the 17 protein-coding genes that were identified as part of the core genome by Oyedara and colleagues but not by our analysis, four were missing only in NC01. The remaining 13 were missing in some of the other *Bdellovibrio* genomes according to our analysis, likely due to differences in the parameters, such as alignment length, used by the two software programs to cluster genes into ortholog groups.

To further investigate the four protein-coding genes identified as missing only in NC01, we used BLAST to align representative amino acid sequences from the corresponding ortholog groups against the full set of amino acid sequences annotated in NC01 by PGAP and RAST. These analyses confirmed that two of the four genes are missing in NC01. The two genes are predicted to encode a transporter permease (Bd1145/WP_011163662) and a protein with a conserved domain involved in synthesis of cytochrome c oxidase (Bd3223/WP_011165579). The two remaining genes are predicted to encode a hypothetical protein and the flagellar biosynthesis protein FliL. For each of these genes, BLAST alignments yielded evidence of a possible ortholog within the NC01 genome.

In particular, BLASTP alignment showed 78% sequence similarity between the protein-coding gene annotated as a hypothetical protein (Bd1976/WP_011164428) and a 493 aa sequence in NC01 that was annotated by both PGAP (DOE51_09405) and RAST (peg.1859). The NC01 amino acid sequence is 161 aa longer than the representative amino acid sequence and other sequences in the ortholog group due to a stop site difference. It was clustered in an ortholog group by itself in the analyses of PGAP and RAST protein-coding genes, likely due to the alignment length cutoff. Because the NC01 amino acid sequence shares high sequence similarity with members of the hypothetical protein ortholog group over two-thirds of its length, it may have derived from the same ancestral gene.

Finally, the flagellar biosynthesis protein FliL was identified as missing only in NC01 because it was annotated as a pseudogene by PGAP (DOE51_16425). However, RAST annotated it as a functional protein-coding gene (peg.3231). BLASTP alignment showed 89% sequence similarity between a representative FliL sequence (Bd3329/WP_011165675) and the NC01 amino acid sequence annotated by RAST, but over only 69% of the representative sequence. This is due to differences in predicted start sites and therefore length between the NC01 amino acid sequence (177 aa) and the representative amino acid sequence (236 aa). TBLASTN alignment of the representative amino acid sequence against the region of the NC01 nucleotide sequence annotated as a pseudogene by PGAP (3428266-3428813) confirmed that there is no alternative start site in this region, and the NC01 protein-coding gene is shorter. These analyses explain why the flagellar biosynthesis protein FliL was annotated as a pseudogene by PGAP and therefore not included in the ortholog group generated using PGAP annotations. However, the NC01 protein-coding gene was included in the FliL ortholog group when using RAST annotations.

### Comparative genomics identifies eight regions of unique gene content possibly acquired by *Bdellovibrio* sp. NC01 via horizontal gene transfer

Using the BLAST Ring Image Generator (BRIG), we conducted pairwise comparisons of NC01 to HD100 and W based on nucleotide sequence alignments with BLASTN. We chose HD100 and W based on the 16S rRNA gene phylogenetic tree and average AAI comparisons to represent phylogenetic diversity and genome variation among sequenced intraperiplasmic *Bdellovibrio*. BRIG analysis identified eight NC01 genomic regions of at least 15,000 nucleotides in which no alignments of >200 nucleotides and e-value <0.1 to HD100 or W were reported. These eight regions, which may represent unique NC01 gene content, are marked on Figure 3. The plot of GC content shows that four of the eight regions (regions 1, 4, 5, and 7) encompass substantial decreases in GC content compared to the typical variation along the genome. Such deviation in GC content may be an indicator of horizontal gene transfer (HGT) (56).

**Figure 3.**
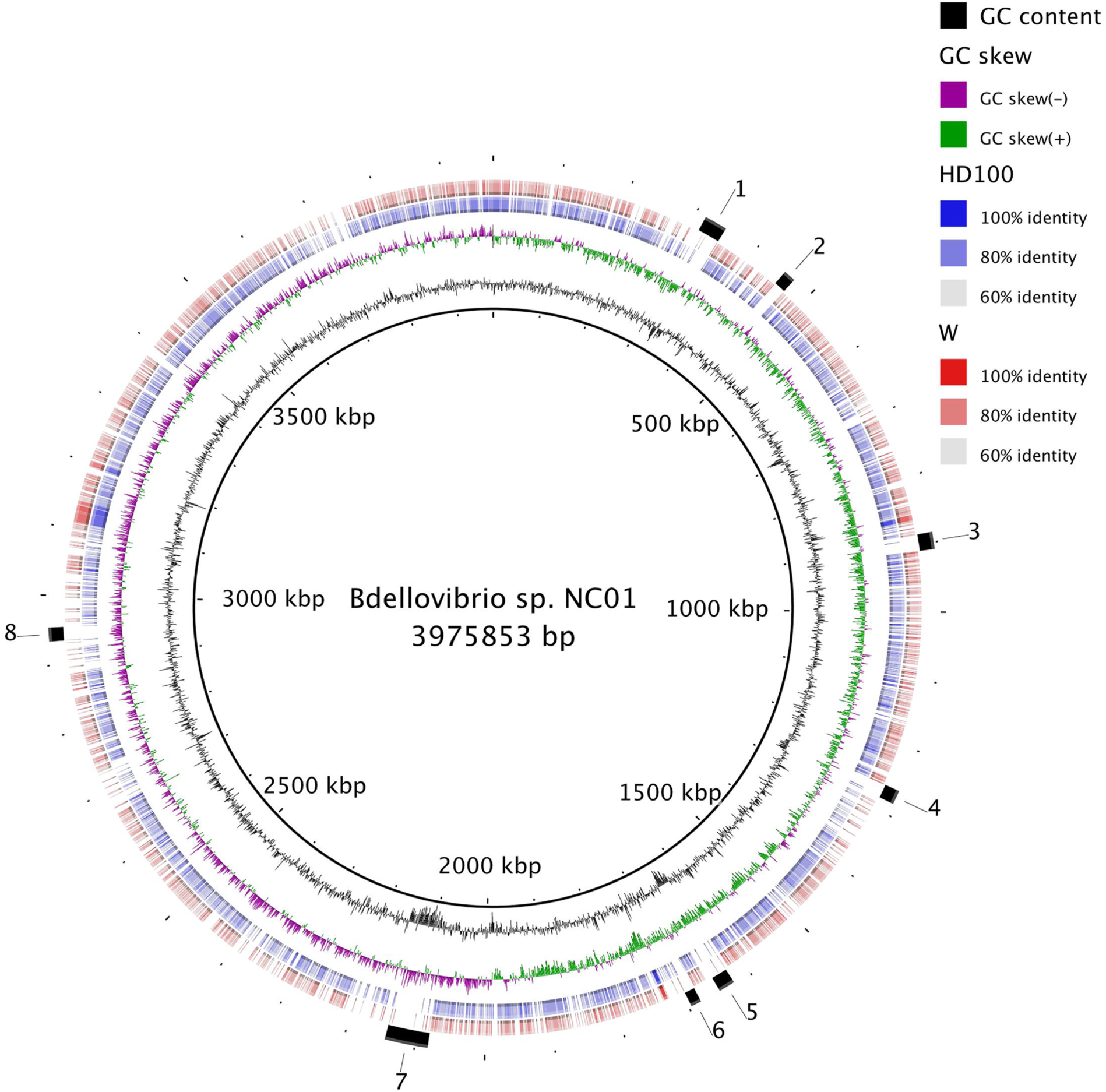
Genome-wide pairwise comparisons of *Bdellovibrio* nucleotide sequences. The genome of *Bdellovibrio* sp. NC01 is shown as the innermost ring. The next ring is a plot of GC content and the purple and green ring is a plot of GC skew, both of which were generated by the BLAST Ring Image Generator (BRIG) using sliding windows to determine deviation from the average value for the genome. The blue ring and red ring show pairwise nucleotide sequence alignments of NC01 to HD100 and W, respectively. Each individual alignment is shown as a block along the ring, and color gradation of the blocks reflects the percent similarity of alignments. The outermost ring denotes eight regions lacking alignments, which may indicate unique NC01 gene content.

To assess whether the genes encoded in these eight NC01 genomic regions are present but divergent in HD100 and W, we used TBLASTN to compare NC01 amino acid sequences annotated in each region by RAST to the HD100 and W genomes translated into all six reading frames. RAST annotated 11 to 50 protein-coding sequences within each of the eight NC01 genomic regions. By searching for these sequences along the entire HD100 and W genomes, this analysis detects similar protein-coding sequences even if the genes have undergone rearrangement. For the majority of the protein-coding sequences in each of the eight NC01 regions, TBLASTN did not produce any alignments to HD100 or W with >70% query coverage. For those protein-coding sequences that did align to HD100 and/or W with >70% query coverage, identity was low (<40%) for most of the sequences. Less than five protein-coding sequences in each region aligned to HD100 or W at >70% coverage and >40% identity. This suggests that the protein-coding sequences in the eight NC01 genomic regions are missing from HD100 and W, instead of divergent.

In addition to comparisons against HD100 and W, we also used BLASTP to align amino acid sequences annotated in each region by RAST against the non-redundant GenBank database. For each sequence, we examined the top 250 alignments for any sequences identified as *Bdellovibrio*. This analysis expands the search space beyond complete *Bdellovibrio* genomes to any deposited sequences that were denoted by the submitters as *Bdellovibrio*. In two of the regions (regions 1 and 4), only one-third of amino acid sequences aligned to *Bdellovibrio* sequences. In every other region except region 8, 52-64% of the amino acid sequences aligned to *Bdellovibrio* sequences. In region 8, three-fourths of the amino acid sequences aligned to *Bdellovibrio* sequences. These results suggest that most of the protein-coding sequences in regions 1 and 4 may be unique to *Bdellovibrio* sp. NC01, because they do not align to other *Bdellovibrio* sequences in the database. In the other regions, the larger proportion of shared gene content between NC01 and *Bdellovibrio* in the database may be the result of HGT events that occurred before the divergence of NC01 and these strains.

The hypothesis that the eight NC01 genomic regions were acquired by horizontal gene transfer is also supported by predicted functions of the protein-coding sequences encoded within the regions. RAST annotation identified functions related to mobile elements in five of the eight regions. These include phage tail fibers (regions 1, 5 and 6), site-specific recombinases or integrases (regions 1, 4, and 7), a phage-related DNA primase (region 4), and a plasmid conjugative transfer protein (region 7). In addition, Prophinder analysis of the complete NC01 genome identified a portion of region 5 (1,632,099 – 1,632,833) as a possible prophage. Three regions (regions 2, 3 and 8) did not encode any obvious mobile element functions.

Apart from mobile element-related genes, most of the protein-coding sequences in the eight NC01 genomic regions are annotated as hypothetical proteins by RAST, therefore it is unclear what functional capacities may have been gained by *Bdellovibrio* sp. NC01 via HGT and whether those functions impact predation. Some of the genes were annotated with predicted functions, which enabled us to assess gene content in the eight regions in terms of broad functional categories. Four of the eight regions (regions 1, 3, 6 and 8) encode predicted functions related to transcriptional regulation, and five regions (regions 1, 3, 5, 6, and 7) encode predicted functions related to transport. Regarding enzymatic activity, region 6 encodes a predicted cell wall endopeptidase, which is interesting given *Bdellovibrio*’s interactions with the Gram-negative prey cell wall during its life cycle.

### Differential gene content between *Bdellovibrio* sp. NC01 and HD100 includes protein-coding genes that may be important for predation

In a pairwise comparison of protein-coding gene content between NC01 and HD100, we found that ~32% of NC01 protein-coding genes are not found in HD100 and ~27% of HD100 protein-coding genes are not found in NC01 (Table 3). To assess broad functional classification of these differentially present genes, we used RAST annotations of the genomes, primarily because, in our experience, RAST predicts functions for more protein-coding genes, whereas PGAP classifies more protein-coding genes as “hypothetical proteins”. Additionally, both genomes were annotated using the same version of RAST, which ensures consistency in gene calls and functional predictions. Of the 1,213 genes identified as present in NC01 and absent in HD100, 516 were assigned a predicted function by RAST, and 697 were annotated as hypothetical proteins. Of the 1,026 genes identified as present in HD100 and absent in NC01, 465 were assigned a predicted function by RAST, and 561 were annotated as hypothetical proteins. Protein-coding genes annotated as “hypothetical proteins” may impact predation phenotype differences between NC01 and HD100; however, the lack of predicted function precludes any hypotheses about the roles of the gene products in predation.

**Table 3.** Differentially present protein-coding genes between *Bdellovibrio* sp. NC01 and *B. bacteriovorus* HD100

|  | PGAP | RAST |
| --- | --- | --- |
| Present in NC01, absent in HD100 | 1175 (31.1%) | 1213 (32.1%) |
| Present in HD100, absent in NC01 | 940 (26.5%) | 1026 (28.5%) |

For the differentially present protein-coding genes assigned a predicted function by RAST, we used InterProScan (57) to assess how the amino acid sequences were grouped into families based on functional or structural classifications. Gene Ontology (GO) classifications are often used for this purpose; however, only half of the amino acid sequences were assigned GO terms. By contrast, Pfam classifications were assigned to 461 of the amino acid sequences identified as present in NC01 and absent in HD100 (89.3% of those annotated with a predicted function) and 408 of the amino acid sequences identified as present in HD100 and absent in NC01 (87.7% of those annotated with a predicted function). Overall, differentially present amino acid sequences belonged to a range of Pfam families, with 349 unique functional or structural classifications for those present in NC01 and absent in HD100 and 336 unique functional or structural classifications for those present in HD100 and absent in NC01.

Previous studies of *B. bacteriovorus* have analyzed gene expression and constructed mutant libraries to identify protein-coding genes likely important for predation. Lambert and colleagues identified 240 protein-coding genes that were up-regulated in HD100 after 30 minutes of interaction between *Bdellovibrio* and *E. coli* prey (50), whereas Duncan and colleagues used a library of *B. bacteriovorus* 109J mutants generated by transposon insertions to identify 104 protein-coding genes that are important for *Bdellovibrio* fitness when preying on planktonic *E. coli* (51). One protein-coding gene (Bd1399) was identified by both studies, yielding a combined list of 343 unique protein-coding genes implicated in successful predation of *E. coli* by *Bdellovibrio*. To determine whether these genes are present in NC01, we used NCBI annotations of the genomes, which allowed us to match the HD100 locus tags referenced in these studies to NCBI amino acid sequence accession numbers used in the identification of ortholog groups.

Of the 343 HD100 protein-coding genes implicated as important for predation on *E. coli*, 90 are missing in NC01 based on lack of a NC01 amino acid sequence in the ortholog group including the HD100 amino acid sequence. This includes 80 from the list of genes identified by Lambert and colleagues and 11 from the list identified by Duncan and colleagues, with Bd1399 included in both sets. Most of these differentially present genes are annotated as hypothetical proteins or proteins of unknown function. However, 14 are annotated with a predicted function. Ten of these may play important roles in predation, including four protein-coding genes predicted to function in peptidoglycan metabolism (Bd0816/WP_011163367: D-alanyl-D-alanine carboxypeptidase, Bd1358/WP_011163852: peptidoglycan-binding protein, Bd1673/WP_011164147: LTA synthase family protein, Bd3575/WP_011165900: lytic transglycosylase domain-containing protein), four annotated as serine proteases (Bd0629/WP_011163194: trypsin-like serine protease, Bd2332/WP_011164755: serine protease, Bd2334/WP_011164757: serine protease, Bd3238/WP_011165592: serine protease precursor), and two protein-coding genes predicted to function in lipid metabolism or transport (Bd0976/WP_011163513: helicase with a phospholipase domain, Bd3162/WP_011165523: kinase). These ten genes may be involved in lytic activity related to the *Bdellovibrio* predatory lifecycle. Our analysis indicates that these genes are missing from NC01, although it is possible that the functions are fulfilled by other protein-coding genes that do not belong to the same ortholog groups as the HD100 protein-coding genes.

### *Bdellovibrio* sp. NC01 kills fewer tested prey strains than HD100 and is less efficient at killing *E. coli* ML35

To assess variation in predatory phenotypes between *Bdellovibrio* sp. NC01 and *B. bacteriovorus* HD100, we assayed prey range differences and predation efficiency. To determine differences in prey range, we tested NC01 and HD100 against a panel of eight Gram-negative bacteria isolated from different environments, including a freshwater stream and soil from a residential area (Table 4). Plaque formation on a bacterial lawn indicates that the predatory isolate is able to attack and lyse that particular Gram-negative prey strain. NC01 formed plaques on lawns of five of the eight prey strains tested, whereas HD100 formed plaques on lawns of all eight prey strains. NC01 was not able to attack and lyse two prey strains belonging to *Acinetobacter* and *Raoultella* that were isolated from a freshwater stream. In addition, although NC01 formed plaques on *E. coli* ML35, it did not attack and lyse *E. coli* 0057.

**Table 4.** Prey range differences between *B. bacteriovorus* HD100 and *Bdellovibrio* sp. NC01

| Genus | Strain ID | Environment | Plaque formation<sup>a</sup> |  |
| --- | --- | --- | --- | --- |
|  |  |  | HD100 | NC01 |
| *Acinetobacter* | 0036 | Freshwater | Yes | No |
| *Aeromonas* | 0031 | Freshwater | Yes | Yes |
| Enterobacteriaceae | 0032 | Freshwater | Yes | Yes |
| *Raoultella* | 0037 | Freshwater | Yes | No |
| *Pseudomonas* | NC02 (0042)<sup>b</sup> | Soil | Yes | Yes |
| *Serratia* | 0043 | Soil | Yes | Yes |
| *Escherichia* | 0057 |  | Yes | No |
| *Escherichia* | ML35 |  | Yes | Yes |

<sup>a</sup>Plaque formation data based on three replicates
<sup>b</sup>*Pseudomonas* sp. NC02 is referred to as 0042 in (24)

To measure predation efficiency, we quantified viable *E. coli* ML35 by calculating cfu/ml at different time points over 72 hours of co-culture with NC01 or HD100 (Figure 4). In the absence of predatory bacteria, we did not observe any reduction in viable *E. coli* ML35, demonstrating that co-culture conditions do not adversely affect the ML35 population. After 6 hours of co-culture, NC01 did not appreciably reduce viable *E. coli* ML35 compared to the control, whereas HD100 reduced viable ML35 by approximately 1 log unit. After 24 hours of co-culture, there was a substantial difference in the impact of NC01 and HD100 on the *E. coli* population. NC01 reduced viable ML35 by approximately 3 log units compared to the control, whereas HD100 reduced viable ML35 by 6 log units. Over the next 48 hours, HD100 continued to restrict the viable ML35 population, but the population recovered in co-cultures with NC01, increasing by 2 log units. ML35 population recovery in the presence of NC01 was consistently observed in all assay replicates, and cfu/ml plates showed no evidence of contaminant bacteria.

**Figure 4.**
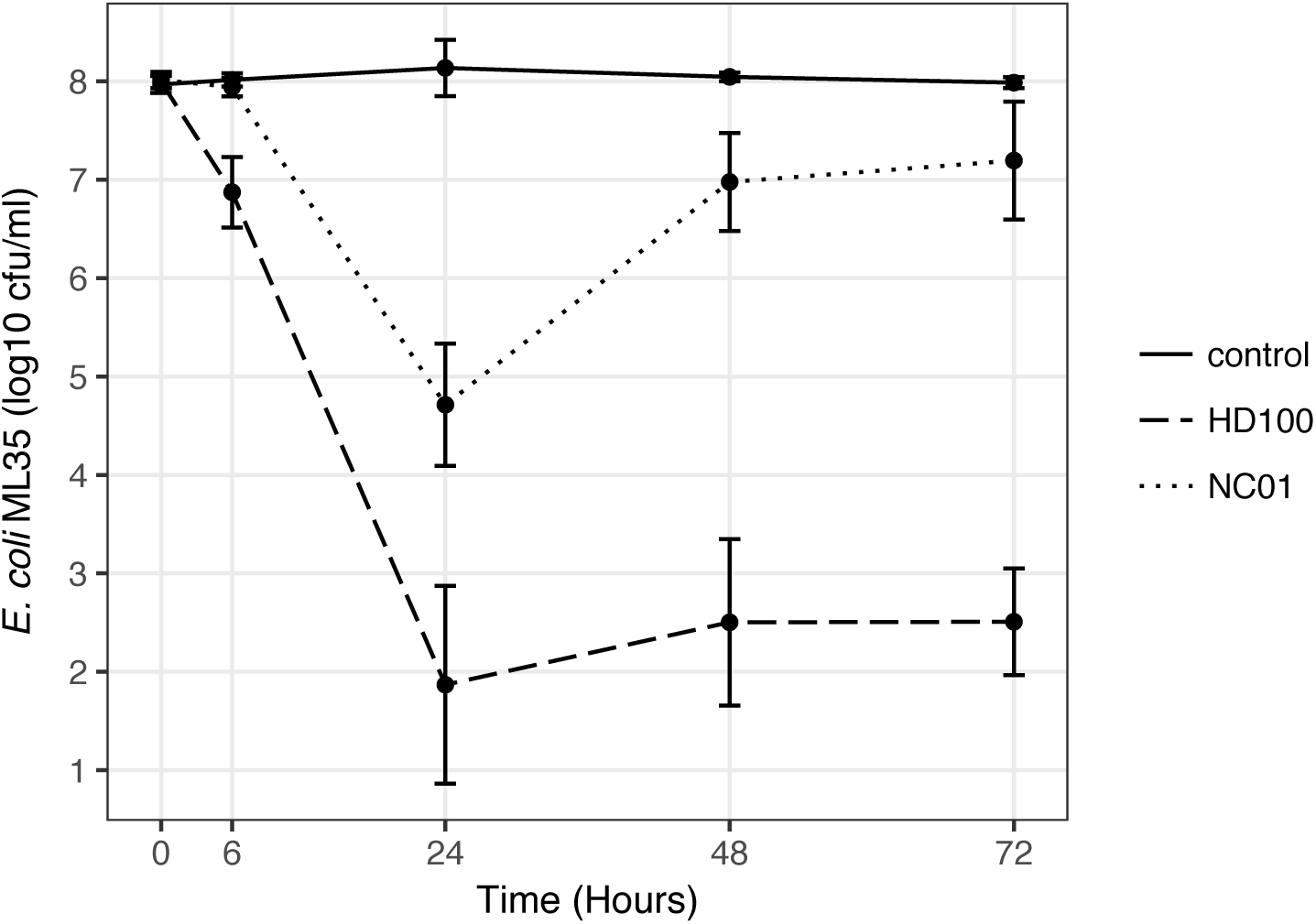
Efficiency of *Bdellovibrio* sp. NC01 and *B. bacteriovorus* HD100 predation on *E. coli* ML35. We quantified viable *E. coli* ML35 over 72 hours on its own (control), combined with *Bdellovibrio* sp. NC01 (NC01) and combined with *B. bacteriovorus* HD100 (HD100). The resulting cfu/ml data were log transformed and plotted using ggplot in RStudio. Each time point shows the mean and standard deviation for at least five replicates.

We quantified NC01 and HD100 after 72 hours of co-culture with *E. coli* ML35 to determine if recovery of the ML35 population is associated with changes in predatory bacteria population density. Using direct cell counts to estimate *Bdellovibrio* cells/ml at the start of co-culture and plaque counts to estimate pfu/ml after 72 hours of co-culture, we observed an increase in the HD100 population of 1 log unit, whereas the NC01 population decreased by 1 log unit (Figure 5). When compared in validation tests, direct cell counts and plaque counts yielded comparable results (Supplementary Figure 1), therefore the observed decrease in NC01 population is not due to a difference in methods for determining predator population density. These data demonstrate that for both NC01 and HD100, active predatory bacteria are present after 72 hours of co-culture with *E. coli* ML35, but, in contrast to HD100, the NC01 population density decreases over the course of the predation efficiency assay.

**Figure 5.**
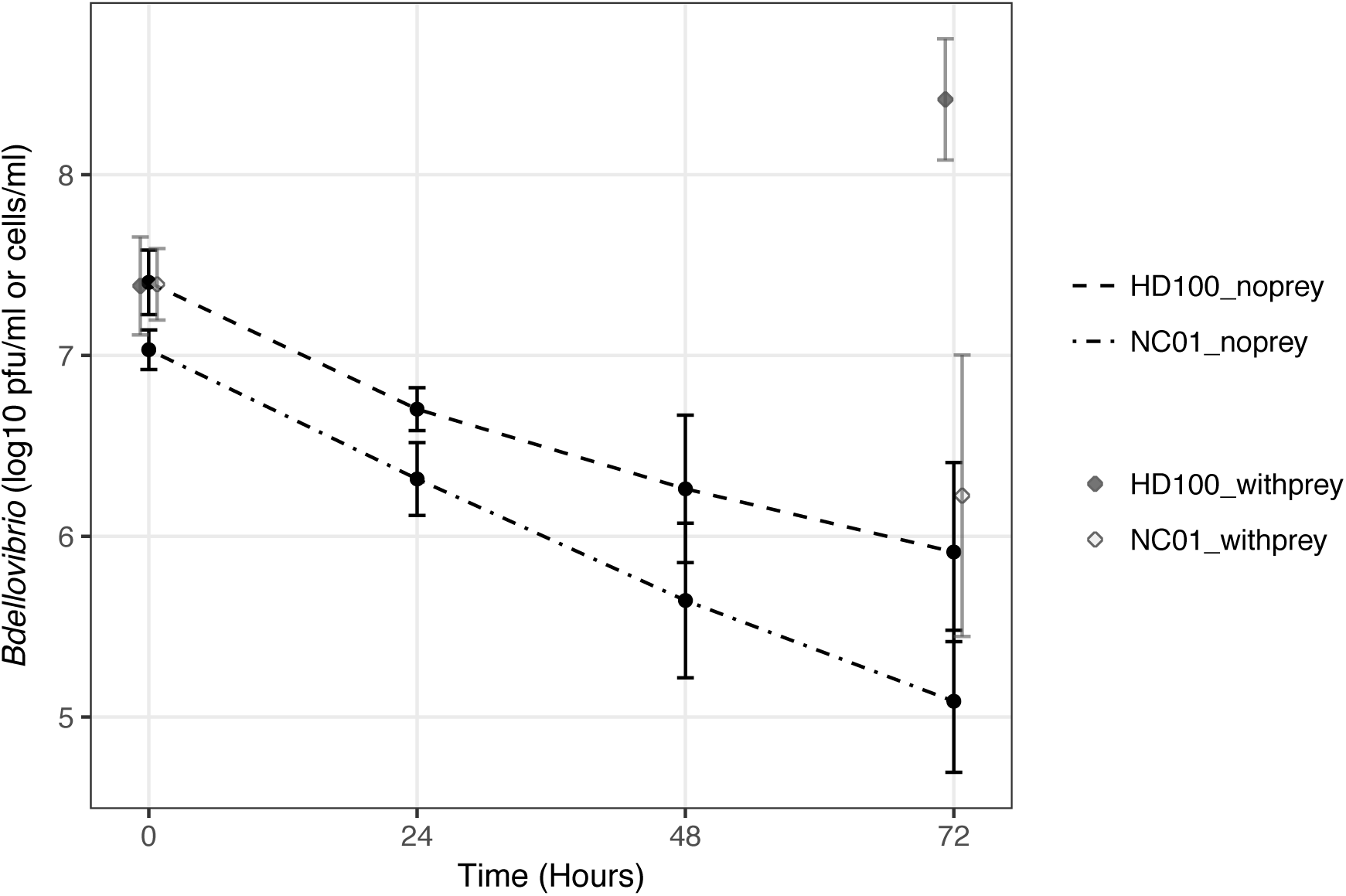
Population density of *Bdellovibrio* sp. NC01 and *B. bacteriovorus* HD100 in the presence and absence of *E. coli* ML35. We quantified *Bdellovibrio* cells/ml at the start of the predation efficiency assay and *Bdellovibrio* pfu/ml after 72 hours of co-culture with *E. coli* ML35 (diamonds). For comparison, we quantified *Bdellovibrio* pfu/ml over 72 hours of incubation in HM buffer alone (circles). The resulting cells/ml and pfu/ml data were log transformed and plotted using ggplot in RStudio. Each time point shows the mean and standard deviation for all replicates.

To determine how predator population density during co-culture with *E. coli* ML35 compares to *Bdellovibrio* survival without prey, we quantified NC01 and HD100 over 72 hours of incubation in HM buffer alone. From similar starting concentrations, the viable NC01 and HD100 populations both decreased by 2 and 1.5 log units, respectively. Overall, the combined data on *Bdellovibrio* population density in the presence and absence of prey show that NC01 population density is higher in co-culture with *E. coli* ML35 than in the absence of prey, which indicates that some NC01 attack phase cells are successfully invading ML35 prey cells and replicating to produce progeny. However, the data also show that the decrease in the NC01 population over the course of the predation efficiency assay is similar to that experienced by both NC01 and HD100 in starvation conditions. The death of NC01 cells and lack of production of new NC01 progeny occurs in the predation efficiency assay despite the presence of viable *E. coli* ML35.

### Plaques formed by *Bdellovibrio* sp. NC01 on *E. coli* ML35 prey lawns are not as clear as those formed by HD100

In addition to quantitative differences in the ability of *Bdellovibrio* sp. NC01 and *B. bacteriovorus* HD100 to kill *E. coli* ML35, we also observed qualitative differences in the appearance of plaques formed by each strain on lawns of ML35 (Figure 6). In contrast to HD100, which forms completely clear, regular plaques, NC01 forms plaques of a similar size, but with incomplete clearance of the prey lawn within the plaques. Some NC01 plaques remain completely turbid over eight days of incubation, whereas others retain a turbid center but begin to clear on the edges of the plaque.

**Figure 6.**
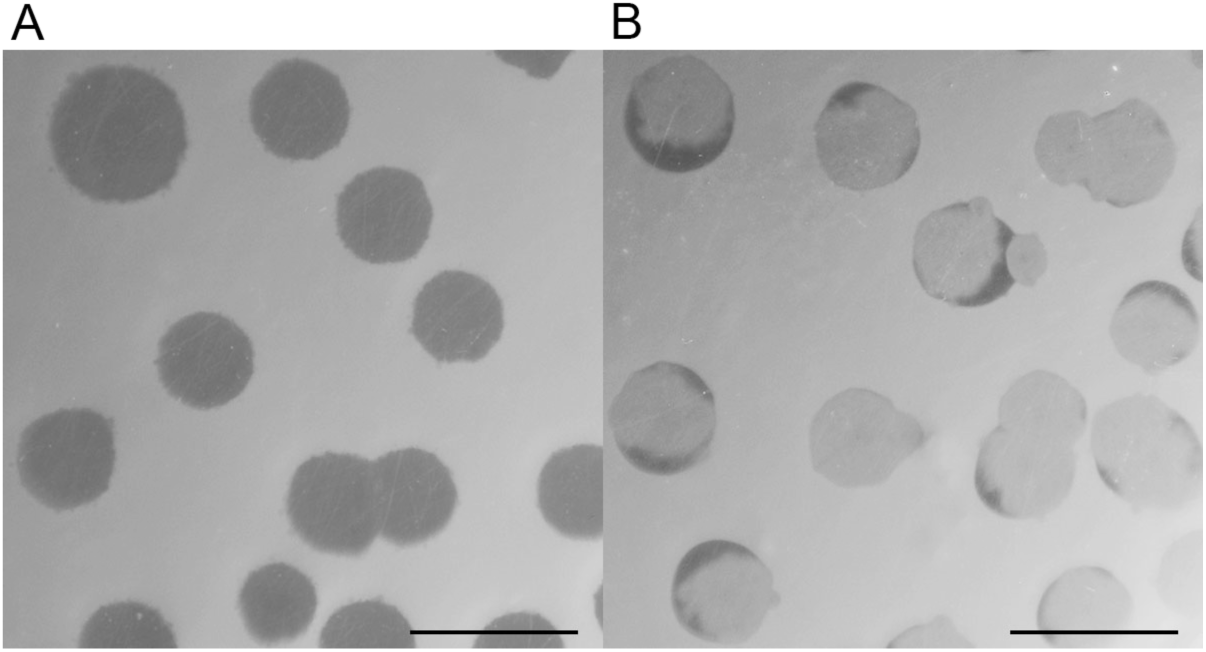
Plaque phenotypes of *Bdellovibrio* sp. NC01 and *B. bacteriovorus* HD100. Using a double agar overlay method, we assessed plaque formation on lawns of *E. coli* ML35. These images were taken after 6 days of incubation and illustrate reproducible differences in the appearance of plaques formed by HD100 (A) and NC01 (B). The scale bar is 10 mm.

## Discussion

We isolated *Bdellovibrio* from soil collected at a manmade landscape feature designed for stormwater management on a college campus. By comparing 16S rRNA gene sequences and analyzing average amino acid identity (AAI) among multiple sequenced *Bdellovibrio*, we determined that the *Bdellovibrio* isolate was sufficiently divergent from *B. bacteriovorus* type strain HD100 and other *B. bacteriovorus* strains that it should be identified as a different species. The only other species currently described within *Bdellovibrio* is *exovorus*; however, this species uses an epibiotic predation strategy that differs from the cell invasion strategy observed by microscopy for the soil *Bdellovibrio* isolate described here. Pending further characterization, we designated the isolate as *Bdellovibrio* sp. NC01.

Because *Bdellovibrio* sp. NC01 is phylogenetically divergent from HD100 and other intraperiplasmic *Bdellovibrio* with sequenced genomes, this isolate provides a valuable opportunity to assess variation in genome content and predatory phenotypes among *Bdellovibrio*. We clustered amino acid sequences from NC01, all five publicly available completely sequenced *Bdellovibrio bacteriovorus*, and *Bdellovibrio exovorus* JSS into ortholog groups. We then compared our analysis with a previous analysis of conserved protein-coding genes reported in (23), which included the same six publicly available *Bdellovibrio* genomes and an additional partially sequenced genome of a *Bdellovibrio* strain isolated from soil. This comparison identified 778 protein-coding genes conserved among all eight *Bdellovibrio* genomes. Intraperiplasmic *Bdellovibrio* genomes range in size from 3 - 4 Mb with 2,760 - 3,780 annotated protein-coding genes, whereas the genome of the epibiotic predator *Bdellovibrio exovorus* JSS is smaller at 2.66 Mb with 2,597 annotated protein-coding genes. Our analysis of the *Bdellovibrio* “core genome” indicates that approximately 20-30% of the protein-coding genes encoded by a given *Bdellovibrio* genome are conserved among intraperiplasmic and epibiotic *Bdellovibrio*.

Analysis of the *Bdellovibrio* core genome also identified four protein-coding genes that appeared to be missing in *Bdellovibrio* sp. NC01 but conserved in the other seven *Bdellovibrio*. We confirmed the absence of two protein-coding genes using manual BLASTP alignments. One encodes a protein of unknown function. The other is annotated as a transporter permease belonging to a conserved domain family that includes AcrB in *E. coli*. AcrB is the inner membrane component of an efflux pump that transports small molecules such as bile salts and some antibiotic compounds out of the cell (58). In addition, AcrB is involved in contact-dependent growth inhibition, in which a bacterial cell attaches to another cell and inhibits its growth. Mutants that do not express functional *acrB* are resistant to growth inhibition (59). Given these functional roles, the absence of the transporter permease in *Bdellovibrio* sp. NC01 may impact this strain’s susceptibility to small inhibitory molecules.

For the remaining two protein-coding genes that appeared to be missing only in NC01, manual BLAST alignments provided evidence of an ortholog encoded by the NC01 genome. In particular, the PGAP annotation of NC01 flagged a gene encoding the flagellar biosynthesis protein FliL as a pseudogene; therefore, it was not included in an ortholog group in our analysis. However, RAST annotated this gene as a functional protein-coding gene, although the resulting protein product is 59 aa shorter than other *Bdellovibrio* FliL due to a difference in the location of the start site. Given the shortened length of this NC01 amino acid sequence, it is unclear whether it is a fully functional flagellar biosynthesis protein. Negative stain electron microscopy of NC01 attack phase cells shows a single, polar flagellum; therefore, NC01 is clearly able to assemble flagella. In addition to the gene discussed here, RAST annotated two other protein-coding genes as flagellar biosynthesis protein FliL. All three of the resulting amino acid sequences are classified into the same Pfam family (pfam03748); therefore, the NC01 genome may encode functionally redundant genes.

Genome-wide comparisons of NC01 to HD100 and W using BLASTN identified eight regions of unique gene content along the NC01 genome, and TBLASTN confirmed that the protein-coding sequences within these regions are missing from the genomes of HD100 and W. Multiple lines of evidence support the hypothesis that these eight regions were acquired by *Bdellovibrio* sp. NC01 via horizontal gene transfer. Four of the regions have substantially lower GC content than the genome average, which may indicate horizontal acquisition from a bacterial genome with a lower average GC content than that of NC01 (56). Five of the regions encode genes with predicted functions related to mobile elements such as phage and plasmids, and one of these regions was identified as a possible prophage by phage finding sequence analysis software.

Alignment by BLASTP of amino acid sequences encoded within two of the eight regions against the GenBank database suggested that these two regions may have been acquired recently by *Bdellovibrio* sp. NC01, because most of the sequences did not align to *Bdellovibrio* sequences (complete or partial) in the database. This indicates that, given the current representation of *Bdellovibrio* in the database, gene content in these two regions is mostly unique to NC01. For the other six regions, the majority of amino acid sequences encoded in these regions aligned to *Bdellovibrio* sequences in the database. This suggests that these *Bdellovibrio* strains may be closely related to NC01, and the HGT events corresponding to acquisition of these six regions occurred in the ancestral lineage before NC01 diverged from these strains. Addition of more complete *Bdellovibrio* genomes to the database will improve our ability to reconstruct evolutionary events such as horizontal gene transfer.

The analysis of *Bdellovibrio* sp. NC01 gene content reported here further illustrates the role of horizontal gene transfer as an important force in predatory bacteria genome evolution. Comparative genomics has yielded evidence of both ancient and recent HGT in *Bdellovibrio* and in the saltwater-adapted predatory genus *Halobacteriovorax* (24, 60–62). However, lack of functional annotation of many *Bdellovibrio* protein-coding sequences prevents a thorough assessment of how HGT events impact predatory phenotypes. Within the eight NC01 genomic regions likely acquired via HGT, many of the protein-coding sequences are annotated as hypothetical proteins, which provides no insight into their function and their potential roles in the predatory life cycle. To fully understand how *Bdellovibrio* genome variation relates to variation in predatory phenotypes, it is essential to assign functions to protein-coding genes.

As the type strain, *B. bacteriovorus* HD100 has been the focus of multiple studies investigating molecular mechanisms governing predation. We compared protein-coding gene content between NC01 and HD100 and showed that ~30% of genes found in one of the strains are missing in the other. These differentially present genes encode proteins from a wide range of functional and structural families. To further investigate the possible impact of differentially present genes on predation by NC01, we assembled a set of 343 HD100 protein-coding genes identified as important for predation by previous studies using gene expression analysis or mutagenesis (50, 51). About a quarter of these genes are missing in NC01. Most are annotated as hypothetical proteins, but ten were annotated with functions that may be important for lytic activity at different stages of the predatory lifecycle. In particular, four HD100 protein-coding genes that are missing in NC01 are predicted to play a role in peptidoglycan metabolism. Invasion of a prey cell by *Bdellovibrio* involves opening a hole in the prey cell peptidoglycan, then resealing and reshaping peptidoglycan to form a rounded bdelloplast (11, 15). If NC01 is missing enzymes that play a role in these processes, it may impact the ability of this strain to invade prey cells and establish itself in the prey periplasm.

In addition to genome comparisons, we also compared predatory phenotypes between *Bdellovibrio* sp. NC01 and *B. bacteriovorus* type strain HD100. Based on plaque formation on lawns of eight different Gram-negative bacteria, NC01 has a more restricted prey range than HD100. In particular, HD100 was able to attack and kill both tested strains of *E. coli*, whereas NC01 was only able to attack and kill *E. coli* ML35. Other researchers have also observed that a particular *Bdellovibrio* strain may vary in its ability to kill different members of the same species (17, 63); however, the mechanisms governing these outcomes are not known. By demonstrating that strain-level variation within a Gram-negative species determines susceptibility to predation, these data have potentially important implications for developing predatory bacteria as biocontrol agents in the treatment of bacterial infections. If a *Bdellovibrio* isolate attacks and kills one representative of a Gram-negative species, it does not guarantee that the isolate will attack and kill other strains within the species. Further work to define the molecular mechanisms underlying variation in prey range may enable prediction of whether a Gram-negative strain will be susceptible to predation by a particular *Bdellovibrio* strain.

Although both NC01 and HD100 can attack and kill *E. coli* ML35, predation by HD100 causes a greater reduction in the ML35 population after 24 hours *in vitro* compared to predation by NC01. HD100 then maintains a low ML35 population density, although it does not completely eliminate viable ML35. By contrast, in co-cultures of NC01 and ML35, the ML35 population recovers after 24 hours, whereas the NC01 population decreases. This is likely not the result of general predator-prey dynamics, such as an increase in search time due to reduced ML35 numbers, because we did not observe ML35 population recovery in co-cultures with HD100. In contrast to NC01, the HD100 population increases after co-culture with *E. coli* ML35. When incubated under similar conditions but without prey, both NC01 and HD100 populations decrease. Overall, data on *Bdellovibrio* population density suggest that NC01 is experiencing starvation conditions during co-culture with *E. coli* ML35 despite the presence of viable prey.

Differences in predation efficiency between NC01 and HD100 on *E. coli* ML35 may also be related to variation in plaque phenotype. Plaques formed by NC01 on lawns of ML35 are turbid, whereas plaques formed by HD100 are completely clear. Turbid plaques have been reported as a characteristic of prey-independent mutants of *Bdellovibrio*; however, these plaques are typically smaller than those formed by the wild-type strain and sometimes have a small colony of host-independent *Bdellovibrio* at their center (64, 65). Neither of these features were observed when comparing NC01 and HD100 plaques, suggesting that the NC01 plaque phenotype is not the result of prey-independent growth.

The predatory phenotypes observed during NC01 co-culture with *E. coli* ML35 may be the result of a mechanism specific to the prey, the predator, or a combination of the two. Other researchers have observed prey population recovery (66, 67), and Shemesh and Jurkevitch demonstrated that recovery of *Erwinia carotovora* in co-cultures with *Bdellovibrio* isolated from soil was due to development of phenotypic resistance to predation. It is possible that *E. coli* ML35 uses a similar mechanism to develop resistance to predation by *Bdellovibrio* sp. NC01, thereby causing the NC01 population to decrease as a result of starvation due to lack of susceptible prey. This mechanism is either not expressed or not successful in co-cultures with *B. bacteriovorus* HD100.

It is also possible that *E. coli* ML35 remains susceptible to predation throughout co-culture, but NC01 is deficient in some aspect of the predatory lifecycle. Our comparison of NC01 and HD100 protein-coding gene content indicated that NC01 is missing some lytic enzymes that may be important for predation. If these functions are not fulfilled by other protein-coding genes, multiple stages of the predatory lifecycle may be affected, resulting in reduced predation efficiency compared to HD100. For example, the turbid plaques formed by NC01 on lawns of *E. coli* ML35 may arise from incomplete lysis of ML35 cells. If prey cells are successfully invaded but not lysed, *Bdellovibrio* progeny will be trapped, which would explain the incomplete clearance of the prey lawn and the decrease in the NC01 population observed during predation efficiency assays. Explanations of NC01 predatory phenotypes that are specific to prey, such as plastic phenotype resistance, and those specific to NC01 itself, such as deficient or missing lytic enzymes, may not be mutually exclusive. Further work will test these explanations.

Overall, comparisons between *Bdellovibrio* sp. NC01 and *B. bacteriovorus* HD100 identified variation in predation phenotypes that may have important implications for the ecological impact of each of these strains and their effectiveness in clinical applications. The molecular mechanisms governing phenotypes such as prey range and predation efficiency are not known. Comparative genomics can generate hypotheses linking gene content differences to observed phenotype differences, but the lack of functional annotation for many protein-coding genes complicates this effort. It is also likely that the outcomes of interactions between predatory bacteria and prey, such as whether or not two different *Bdellovibrio* can kill the same Gram-negative strain, are dependent upon mechanisms specific to the *Bdellovibrio* strain and mechanisms specific to the prey strain. Defining these mechanisms is key for understanding variation in predatory phenotypes among divergent *Bdellovibrio*.

## Acknowledgements

We thank Lisa Sadzewicz and Luke Tallon at the Institute for Genome Sciences at the University of Maryland Baltimore for PacBio sequencing services and Janet Atoyan at the Rhode Island Genomics and Sequencing Center for Illumina MiSeq sequencing services. We thank Sean O’Donnell for isolating freshwater prey bacteria and Mark Martin and Brett Pellock for donating bacterial strains. We thank Joey Cerra for contributing some of the scripts used for comparative genomics.

---

**Supplementary Figure 1.**
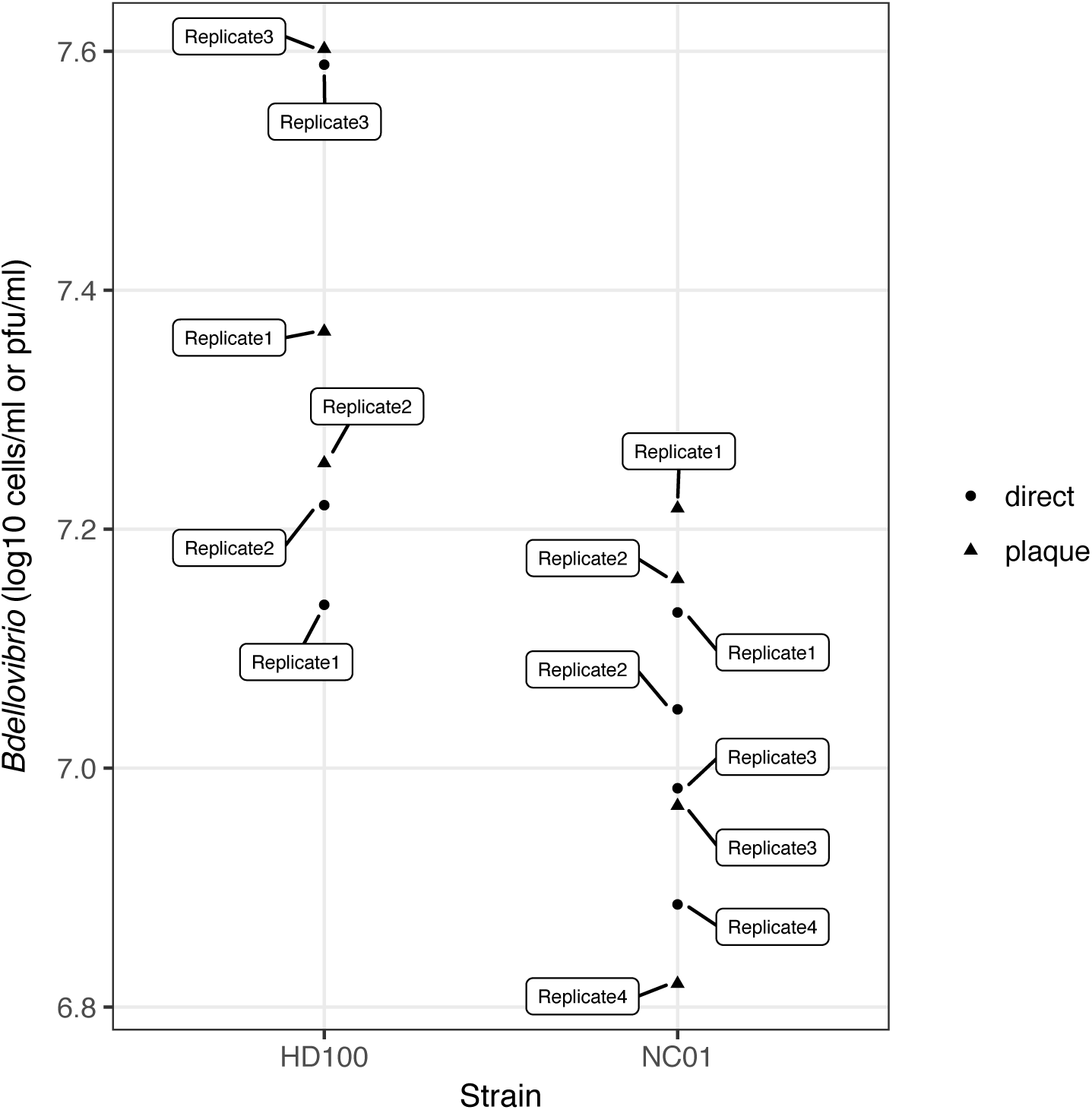
Quantification of *Bdellovibrio* using direct cell counts and plaque counts. For each replicate of the survival assay without prey, initial concentrations of *Bdellovibrio* HD100 and NC01 were estimated by both direct cell counts using a Petroff-Hausser counting chamber (circles) and plaque counts using serial dilution and double agar overlay (triangles). Replicates are labeled.

